# Persistent patterns of high alpha and low beta diversity in tropical parasitic and free-living protists

**DOI:** 10.1101/166892

**Authors:** Guillaume Lentendu, Frédéric Mahé, David Bass, Sonja Rueckert, Thorsten Stoeck, Micah Dunthorn

## Abstract

Animal and plant communities in tropical rainforests are known to have high alpha diversity within forests, but low beta diversity between forests. By contrast, it is unknown if the microbial protists inhabiting the same ecosystems exhibit similar biogeographic patterns. To evaluate the biogeographies of soil protists in three lowland Neotropical rainforests using metabarcoding data, we estimated taxa-area and distance-decay relationships for three large protist taxa and their subtaxa, at both the OTU and phylogenetic levels, with presence-absence and abundance based measures, and compared the estimates to null models. High local alpha and low regional beta diversity patterns were persistently found for both the parasitic Apicomplexa and the free-living Cercozoa and Ciliophora, even though they have different ecological functions and different dispersal modes. In comparison to the null models, both OTU and phylogenetic diversities showed spatial structures between forests, but only phylogenetic diversity showed spatial patterns within forests. These results suggest that the biogeographies of macro-and micro-organismal eukaryotes in lowland Neotropical rainforests are partially structured by the same general processes. As with arthropods, the protists’ high alpha diversity within forests presents problems for estimating their local diversity, and shows that regional diversity cannot be easily estimated because of low turnover between forests.

## Introduction

The distributions of animal and plant species within and between forests has been a central focus in tropical biology research (Connell 1978, Hubbell 2001). Understanding these macro-organismal biogeographic patterns has enabled biologists to establish conservation strategies (Hubbell et al. 2008, Moritz et al. 2001, Turner and Corlett 1996), to infer how diversity arose through the combination of historical, evolutionary and ecological processes (Chave et al. 2002, Myers et al. 2013, Rosindell et al. 2011), and to estimate the total diversity of different taxa (Basset et al. 2012, Novotny et al. 2007, Slik et al. 2015). One way of looking at these patterns is to evaluate taxa-area relationships (TAR), which measures the increase in richness as more area is sampled (Arrhenius 1921, Drakare et al. 2006). Another way is to evaluate distance-decay relationships (DDR), which measures the decrease of community similarity in relation to distance (Morlon et al. 2008, Soininen et al. 2007). Together these patterns allow for comparing the biogeography of taxa that have different ecological functions and different dispersal modes (Novotny et al. 2007, Saito et al. 2015, Wetzel et al. 2012, Zinger et al. 2014).

Most studies inferring TAR and DDR of tropical macro-organisms have focused on species diversity using morphological observations, but a few have used molecular approaches. At local scales, TAR and DDR slopes of arthropods and plants were estimated to be steep, reflecting high alpha diversity and high turnover within forests (Basset et al. 2012, Condit et al. 2002, Hubbell 2001, Novotny et al. 2007). By contrast, TAR and DDR slopes at regional scales were low, reflecting the low beta diversity of animals and plants between forests (Condit et al. 1996, Condit et al. 2002, Novotny et al. 2007, Plotkin et al. 2000). Fewer TAR and DDR studies of tropical macro-organisms have analyzed phylogenetic diversity, in which evolutionary relationships are considered, but these studies have likewise found high local alpha and low regional beta diversity patterns (González-Caro et al. 2014, Swenson et al. 2012, Zhang et al. 2013). The phylogenetic-based TAR slopes were observed to be steeper within forests compared to species-based TAR slopes for mammals and trees, supporting a finer scale biogeographic structure at the phylogenetic level (Mazel et al. 2014, Swenson et al. 2013). In both species-and phylogenetic-based studies of tropical macro-organisms, assessing the non-random origin of the biogeographic patterns were used to unravel spatial vs. non-spatial ecological and evolutionary processes (Hardy 2008, Kraft et al. 2011, Zhang et al. 2013).

Recently it was shown that similar to macro-organisms, soil-inhabiting microbial protists were exceedingly species rich with low similarity among samples within and between lowland rainforests in Costa Rica, Panama, and Ecuador (Mahé et al. 2017). Here we evaluated the taxa-area and distance-decay relationships of this protist diversity from those soil samples at both the OTU-and phylogenetic-diversity levels to ask if they exhibit biogeographical patterns, and if those patterns are similar for taxa with different ecological functions and different dispersal modes. Uncovering these answers allowed us to infer general eukaryotic biogeographic patterns in the Neotropical rainforests for both the macro- and micro-organisms, and to gain insights on how protist distributions affect our ability to estimate total local and regional diversity in the tropics.

## Methods

All codes used here, and all numbers used for the figures, can be found in HTML format (**File S1**). Details of sampling and sequencing can be found in Mahé et al. (Mahé et al. 2017). Briefly, soil samples were taken from: La Selva Biological Station, Costa Rica; Barro Colorado Island (BCI), Panama; and Tiputini Biodiversity Station, Ecuador. DNA from every two consecutive samples was combined in equal concentration to reduce costs, resulting in 144 sample sites. DNA was amplified for the V4 region of 18S rRNA with general eukaryotic V4 primers (2010), and sequenced with Illumina MiSeq. Reads were clustered into OTUs with Swarm v2.1.5 (Mahé et al. 2015) using *d*=1 with the fastidious option on. Chimeras were identified and removed with VSEARCH v1.6.0 (Rognes et al. 2016). Reads and OTUs assigned by Mahé et al. (Mahé et al. 2017) using the Protist Ribosomal Reference (PR^2^) database v203 (Guillou et al. 2013) to the Apicomplexa, Cercozoa, and Ciliophora were used for this study (**Dataset S1**). The taxonomy of Apicomplexa was modified to follow Rueckert et al. (Rueckert et al. 2011): Colpodellidae not in the Apicomplexa, and *Cryptosporidium* in the Gregarinasina.

Using the OTUs, richness as well as Sørensen (based on presence-absence) and Bray-Curtis (based on relative abundance) dissimilarity indexes were estimated using Vegan v2.4-3 (Oksanen et al. 2013). Using OTU representatives, multiple sequence alignments were generated with MAFFT v7.221 (Katoh and Standley 2013) using the FFT-NS-i strategy. Maximum Likelihood trees were inferred with RAxML v7.3.0 (Stamatakis 2006) using the GTR+CAT model and hill-climbing algorithm. The phylogenetic species variability index (Helmus et al. 2007) for alpha diversity was estimated using picante v1.6-2 (Kembel et al. 2010) and for beta diversity using the unweighted (based on presence-absence) and weighted (based on relative abundance) UniFrac distances (Lozupone and Knight 2005) as implemented in phyloseq v1.20.0 (McMurdie and Holmes 2013). Phylogenetic variability was preferred to Faith’s PD index because of its independence from OTU richness, making it more suitable for testing for phylogenetic patterns independently from the amount of diversity covered (Helmus et al. 2007, Helmus and Ives 2012).

All diversity indexes were estimated based on 1000 randomly rarefied OTU tables normalized to the sample with the lowest number of reads. To allow for statistical robustness, samples were analyzed if they gather at least 80,000 Apicomplexa reads, or 1,000 Cercozoa or Ciliophora reads (100, 108 and 110 samples, respectively). Subtaxa within the Apicomplexa, Cercozoa, and Ciliophora were also analyzed if they had at least ten OTUs, and were found in at least five samples within each forest, and each sample had at least 100 reads. Taxa-area relationships (TAR) were estimated using the double logarithmic generalization of Arrhenius’ equation (Arrhenius 1921), *log(S_obs_) = log(c) + z × log(A)*; where *S_obs_* is the species richness or phylogenetic variability, *A* the sampled area, *c* a constant and *z* the slope. Distance-decay relationships (DDR) were estimated using the double log transformation of the Nekola and White formula (Nekola and White 1999): *log(S_com_) =log(a) + *β* × log(D)*; where *S_com_* was the community or phylogenetic similarity (1 – dissimilarity index), *D* the geographic distance, *a* a constant and *β* the slope. Analyses of variance were used to test significance of linear regressions and significant difference between linear regression slopes (i.e. between observed and null model slopes, see below) were assessed using analysis of covariance. Regression statistics were calculated in each 1000 rarefied matrices and averaged, with p-value being corrected for multiple comparisons (Benjamini and Hochberg 1995).

Areas for the TAR were estimated in circle of increasing radius (Zinger et al. 2014) ranging from 0.5 to 5 km with a 0.5 km increment within forests with each sample being successively the circle center. For area between forests, the same principle was applied using only the Panama samples as circle center, with additional radius ranges between 475 and 484 km to cover Costa Rica samples and between 1152 and 1159 km for Ecuador samples, incrementing by 1 km inside each range. Only lowland Neotropical rainforests contained within each circle was considered to calculate the areas. For Costa Rica and Panama locations, tropical rainforest coverage was recovered from the 2012 MODIS Land Cover in native resolution (Channan et al. 2014, Friedl et al. 2010), with Panama areas being overlaid with the Barro Colorado Nature Monument Land boundaries. The regional lowland Neotropical rainforest areas were estimated by overlaying the “Evergreen Broadleaf forest” class of the 2012 MODIS Land Cover in 5’ × 5’ resolution with the “Tropical rain forest” class of the 2010 FAO’s Global Ecological Zones (FAO 2015), keeping only the union of both layers clipped to the Neotropics. Areas were determined using the EASE-Grid equal area projection (Brodzik and Knowles 2002). All analyses were conducted in R (R Core Team 2016).

Two null models were constructed in which only the tested parameter was randomized while keeping all other structural features unrelated to the relevant null hypotheses tested (Hardy 2008). Null model 1: the geographic positions of the samples were randomly shuffled inside each forest or among all three forests. This null model randomized each OTU’s geographic range without modifying the OTU richness and community composition of single samples. This procedure erased potential OTU spatial structures and resulted in non-spatial TAR and DDR (Scheiner et al. 2011). The null hypothesis tested was that the sample’s geographic position was independent from the community composition. Null model 1 TAR and DDR regressions were computed based on OTU richness and community similarity indexes. Null model 2: the OTUs were randomly shuffled along the tip of the phylogenetic tree within the entire regional OTU pool or for each forest OTU pools separately (Hardy 2008, Kembel 2009). This null model allowed the phylogenetic relatedness to vary independently from the OTU spatial range, OTU richness and community composition and is suitable to test for phylogenetic structuration in a geographical context (Hardy 2008). Null model 2 TAR and DDR regressions were computed based on phylogenetic variability and phylogenetic similarity indexes.

## Results

### Taxonomic assignments

Although many protist taxa were identified in the soils that were sampled throughout the three Neotropical rainforests and sequenced with a metabarcoding approach (Mahé et al. 2017), here we only evaluated the three most abundant groups: Apicomplexa, 35,134,460 reads and 12,247 OTUs; Cercozoa, 2,404,404 reads and 5,158 OTUs; and Ciliophora, 1,479,001 reads and 1,903 OTUs (Fig. 1). These OTUs can be viewed as estimates of species diversity in molecular environmental studies of microbes (Bik et al. 2012, de Vargas et al. 2015, Mahé et al. 2017).

**Figure 1.**
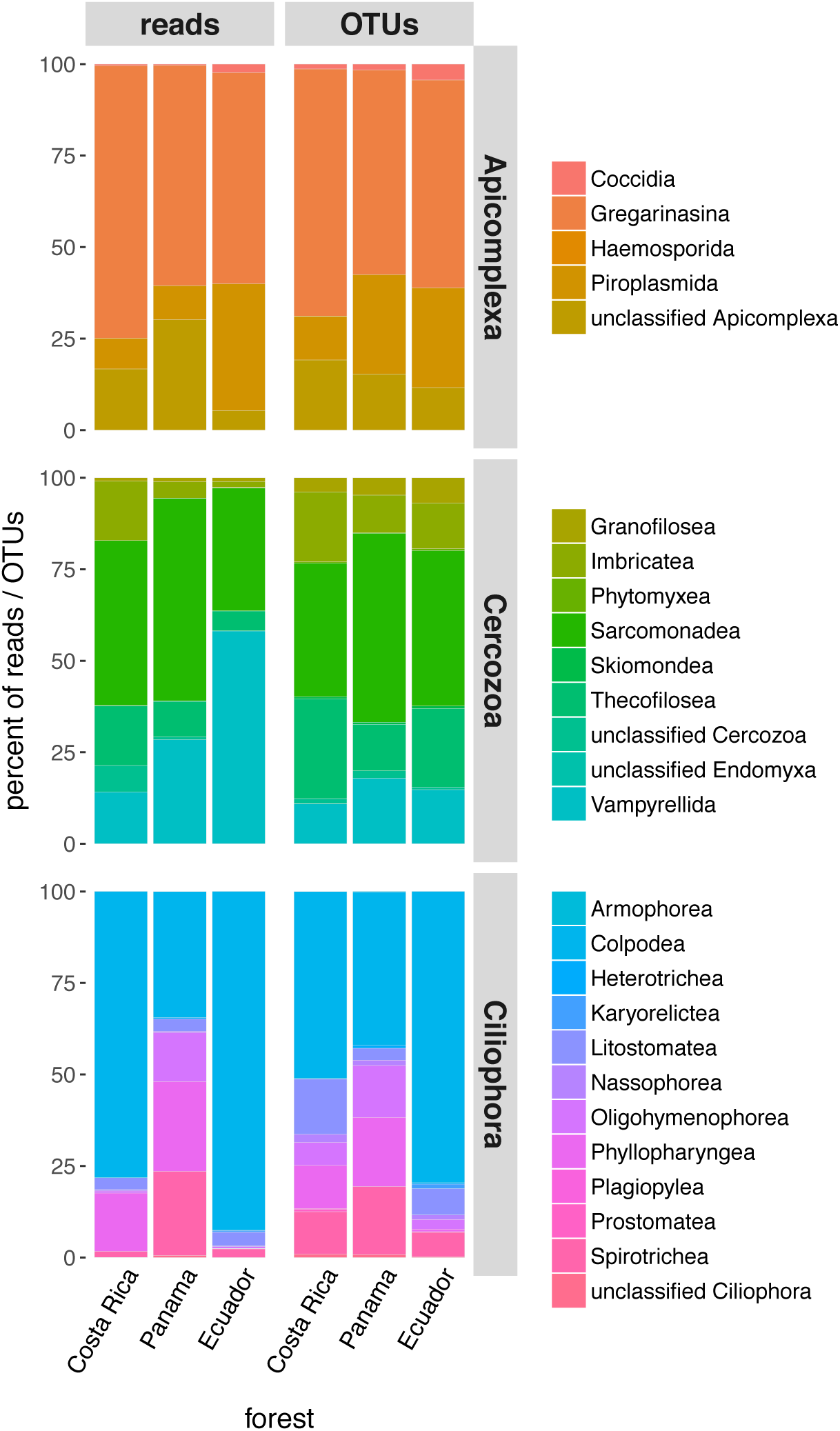
Taxonomic assignment [%] of the reads and OTUs of the parasitic Apicomplexa, and the free-living Cercozoa and Ciliophora, collected in tropical rainforests in Costa Rica, Panama and Ecuador.

Within Apicomplexa, the Gregarinasina, including *Cryptosporidium*, was the dominant taxon (65.9% of the reads, 61.7% of the OTUs). Gregarines are obligate parasites infecting the intestines, coeloms and reproductive vesicles of invertebrates, while *Cryptosporidium* infects the intestines of vertebrates (Desportes and Schrével 2013). Their oocysts and/or gamontocysts are expelled from the macro-organisms with the faeces and can be found in soils worldwide. In *Cryptosporidium*, for example, there are 26 valid species (Ryan et al. 2014), yet we found 4,500 OTUs. The blood parasites in the Haemospororida accounted for only 0.004% of the reads and 0.04% of the OTUs.

The dominant group within Cercozoa was the sarcomonads (44.9% of reads, 38.2% of OTUs), which is comprised of Cercomonadida and Glissomonadida. These two groups are generally bacterivorous or fungivorous flagellates (Bass et al. 2009), although one percent of the glissomonad reads were assigned to an algivorous viridiraptorid genus (Hess and Melkonian 2013). Other detected algivorous cercozoans included the vampyrellid amoebae (Berney et al. 2013). Numerous testate filose amoebae were also found, most of which where the silica-shelled euglyphids and organic-shelled thecofiloseans, but few Endomyxa were detected despite including the plant pathogenic/symbiotic Plasmodiophorida which are often found in other soils (Neuhauser et al. 2014).

Within Ciliophora, Colpodea was the dominant taxon (67.2% of reads, 53.0% of the OTUs). Colpodeans are primarily bacterivores, but some are fungivorous, and have r-selected growth strategies that can quickly dominate the ciliate communities by ex-cysting from their resting stages (Lynn 2008). Although there are fewer than 250 described colpodean species, we found 1,008 OTUs assigned to this taxon. The second and third most abundant ciliates groups were the largely bacterivorous Phyllopharyngea and Spirotrichea. Only one OTU found in Panama was assigned to Armophorea, which thrive in anoxic environments such as insect guts (Lynn 2008).

### Null models

Two null models were computed in order to test for the non-randomness of the below OTU-and phylogenetic-diversity patterns. The null model 1 randomized the sample’s geographic locations inside the considered scale (forest or regional) and was applied to OTU-based diversity measures; this null model allowed the OTU richness accumulation and the OTU composition turnover to change without any spatial constraint (Scheiner et al. 2011). The null model 2 randomized the OTUs along the tips of the phylogenetic trees at the considered scale and was applied to phylogenetic-based diversity measures; this null model allowed phylogenetic relatedness to vary independently from community composition in order to test for sample’s geographic position on phylogenetic diversity only (Hardy 2008).

### Taxa-area relationships

Using OTUs, the TAR slopes (*z*) within forests varied from 0.36 to 0.57 (Fig. 2). These slopes were all significant (p <0.001) with high R^2^ values between 0.71 and 0.84, and none differed significantly from the slopes produced by the null model 1 (ANCOVA p >0.05). TARs between forests (i.e., using all samples from all three forests) were much flatter with slopes from 0.1 to 0.11. These slopes were all significant with low to moderate R^2^ values from 0.25 to 0.60, but all were significantly steeper than null model 1. Subtaxa (i.e., smaller taxa within Apicomplexa, Cercozoa, or Ciliophora) slopes were likewise all significant, with most slopes differing from null model 1 when using all samples (Figs. S2-S3).

**Figure 2.**
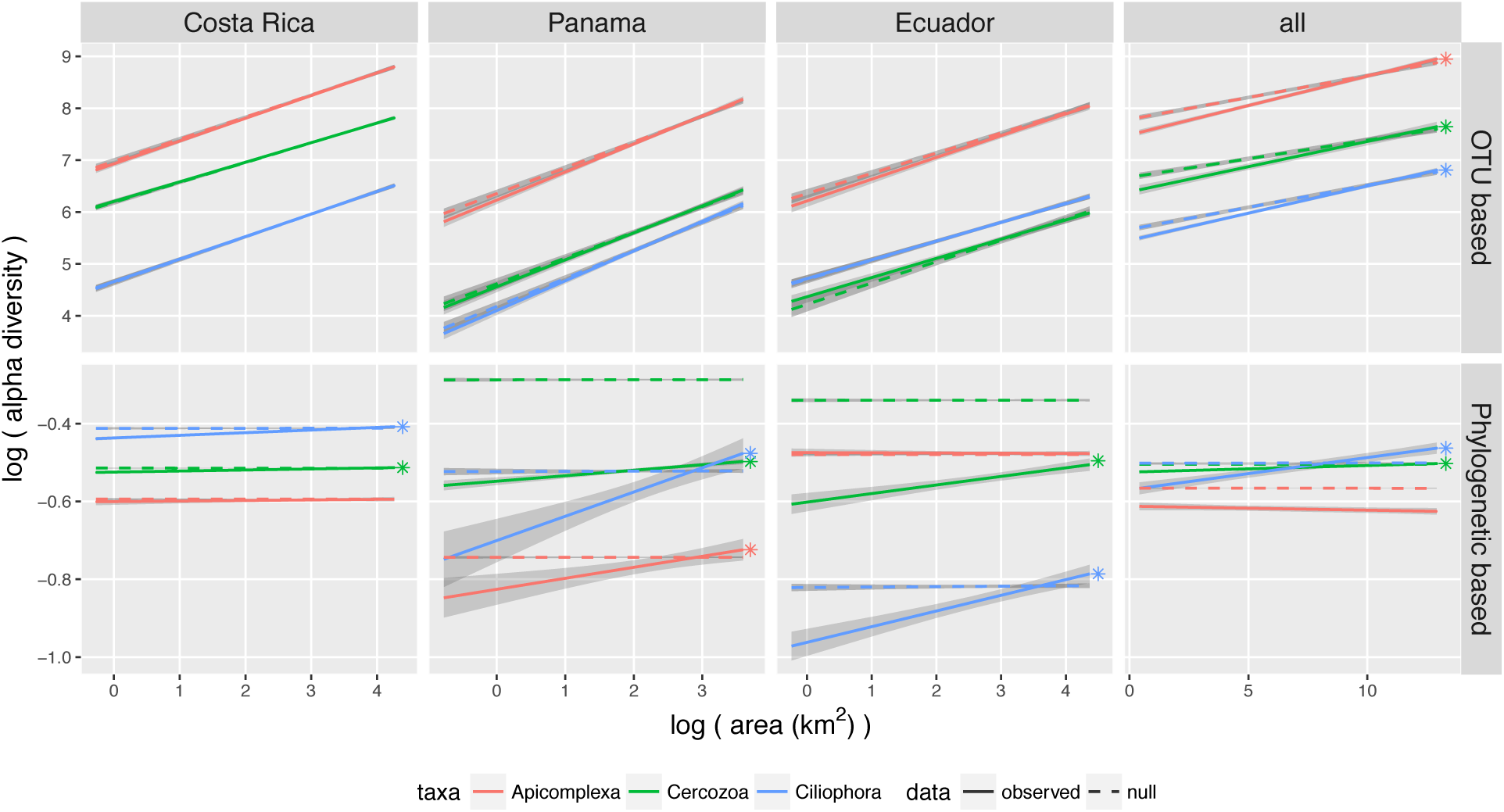
Taxa-area relationships (TAR) within the three forests, and between all sample sites. Shaded areas are 95 % confidence intervals. Slopes were significant for both OTU richness and phylogenetic variability. Slopes were steeper within forest than among forests. Stars denote significant steeper slopes compared to the null model assessed by analyses of co-variance. Null model 1 (random placement of sample locations) was used for OTU based TARs and null model 2 (shuffling of phylogenetic tree tips) for phylogenetic based TARs.

Using phylogenies of the OTUs, the TAR slopes within forests varied from -0.00048 to 0.06 and were mostly significant (Fig. 2). The significant slopes had low R^2^ values from 0.05 to 0.27, and they all differed from null model 2 except in Costa Rica. TARs between forests were much flatter with slopes from -0.001 to 0.0083. Only Cercozoa and Ciliophora had significant slopes that had low R^2^ values from 0.07 to 0.10, but these were significantly steeper than null model 2. Subtaxa slopes were also mostly significantly and mostly differed significantly from null model 2 (Figs. S3-S4).

### Distance-decay relationships

Using OTUs, most DDR slopes (*β*) of protist taxa and subtaxa within forests were not significant using both presence-absence and abundance-based measures, and thus a null model was not evaluated at that scale. DDR slopes of protist taxa between forests varied from -0.31 to -0.22 using presence-absence data, and -0.49 to -0.36 using abundance data (Fig. 3). These slopes were all significant with low R^2^ ranging from 0.05 to 0.28, and all were significantly steeper than null model 1, as the latter resulted in non-significant (i.e., flat) slopes. Subtaxa slopes between forests were all significant and all differed from null model 1 (Fig. S5-S6).

**Figure 3.**
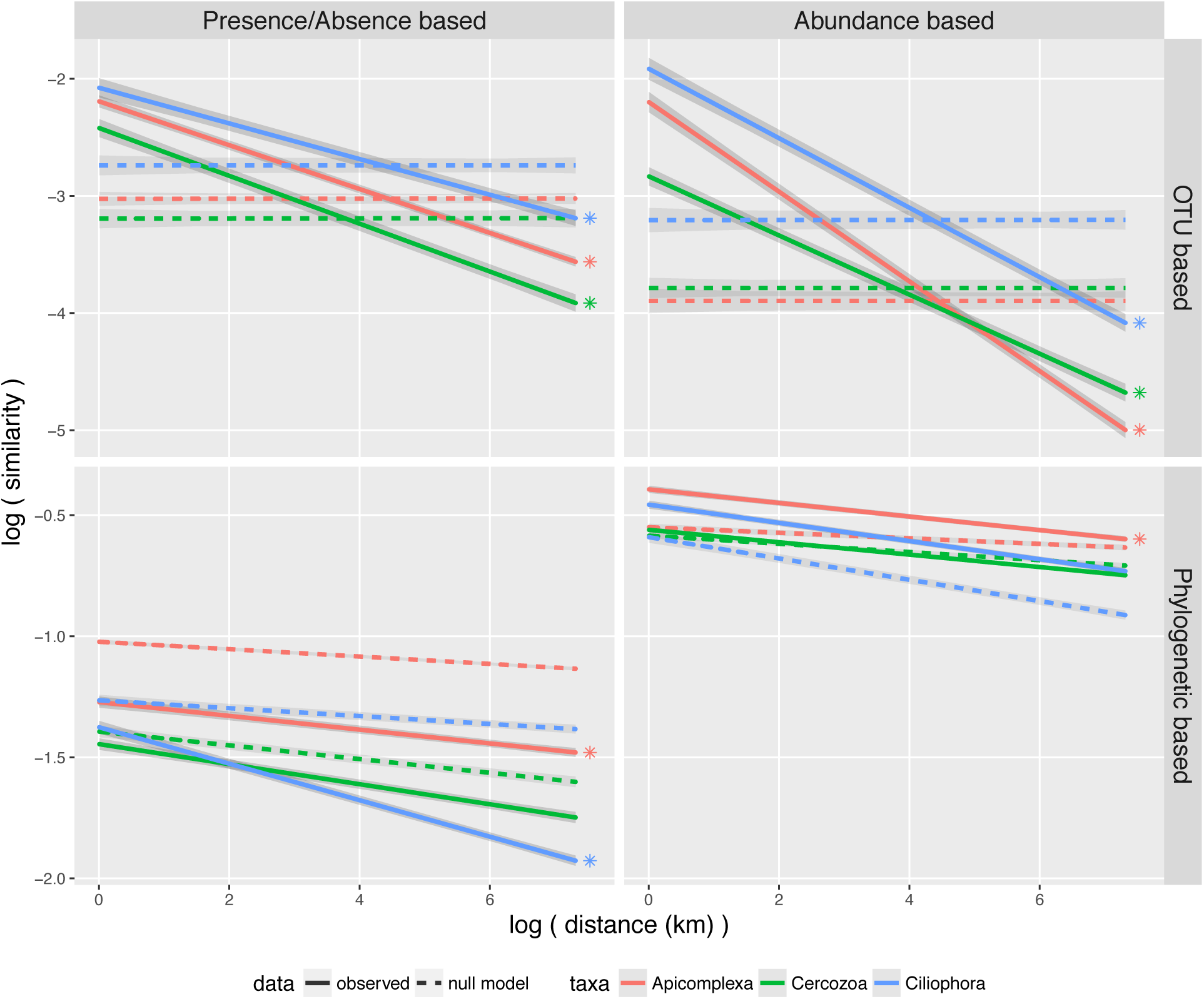
Distance-decay relationships (DDR) for three protist groups using all samples of the three forests. Shaded areas are 95 % confidence intervals. Slopes were not significant within forests for both OTU richness and phylogenetic variability, and therefore are not shown. Stars denote significant steeper slopes compared to null model assessed by analyses of co-variance. Null model 1 (random placement of sample locations) was used for OTU based DDRs and null model 2 (shuffling of phylogenetic tree tips) for phylogenetic based DDRs.

Using phylogenies of the OTUs, most DDR slopes of protist taxa and subtaxa within forests were likewise not significant using both presence-absence and abundance-based measures, and thus a null model was also not evaluated at that scale. DDR slopes of protist taxa between forests varied from -0.08 to -0.03 for presence-absence, and -0.04 to -0.03 for abundance-based (Fig. 3). These slopes were all significant with low R^2^ from 0.04 to 0.16. Only the Apicomplexa for both measures and Ciliophora for presence-absence have significantly steeper DDR slopes than the null model 2. Subtaxa slopes between all samples were mostly significant, but only half differed from null model 2 (Fig. S6).

OTU-and phylogenetic-measures of similarity showed moderate to high correlations within forests (Spearman’s rho from 0.41 to 0.89 for presence-absence, and 0.29 to 0.84 for abundance based measures, p <0.001) as well as when using all samples (from 0.47 to 0.66 for presence-absence, and 0.5 to 0.59 for abundance based measures) (Table S1). These results explain the similar trends between DDR slopes of the two kinds of similarity measures.

## Discussion

At the OTU level, the steep TAR slopes showed that as more area was sampled within a forest there was an increase in the number of observed protists giving rise to high alpha diversity. This increase in OTUs was not due to the accumulation of area *per se*, but due to the accumulation of samples as there was no local difference with the null model; this pattern suggests that non-spatial ecological processes such as environmental filtering drive the within-forest protist diversity. When the area was expanded to include multiple forests the TAR slopes were less steep giving rise to low beta diversity. The steeper regional slopes compared to the null model suggests that spatial ecological processes such as dispersal limitations drive the between-forest protist diversity. The DDR slopes showed that as more samples were taken within a forest there was no significant decrease in community similarity related to distance, but there was a significant non-random decrease when samples were compared between forests. The distance pattern within forests means that no matter how far the samples are from each other they will have few OTUs shared between them, and demonstrates again that protist diversity at the forest scale is not spatially structured. The even lower community similarity between forests also supports the spatial structure of protist community at regional scale.

At the phylogenetic level, TAR and DDR slopes were similar to the OTU level within and between forests, although additional non-random phylogenetic patterns were detected within forests. The area patterns mean that the more area you sample the more new clades you encounter, highlighting a phylogenetic over-clustering of protists at local and regional scales. The absence of geographic structure in phylogenetic similarity within forests reflected the absence of geographic structure in OTU community similarity, as both type of measures were significantly correlated. The distance patterns between forests suggest that, like OTU diversity, phylogenetic diversity is geographically structured at the regional scale. This finding that additional forests contained new OTUs as well as new clades of protists is applicable for the conservation of microbial diversity (Cotterill et al. 2013, Gómez and Nichols 2013, Griffith 2012), as the destruction of unconserved tropical forests would lead not only to the loss of macro-organisms but also to the possible loss of protist clades.

These TAR and DDR patterns of the protist were inferred with our sampling and molecular approaches. The soils were sampled in noncontiguous forests in three countries, but the areas were calculated following a nested sampling scheme (Scheiner 2003). Similar strategies of estimating biogeographic patterns from noncontiguous sampling were used in previous studies of aquatic and terrestrial macro-and micro-organisms (Bates et al. 2013, Martiny et al. 2011, Mazel et al. 2014, Novotny et al. 2007, Venter et al. 2017, Zinger et al. 2014). The molecular data were clustered into OTUs with Swarm, which can be more accurate than other clustering methods that rely on global clustering thresholds (Forster et al. 2016, Mahé et al. 2015). Like any clustering method, Swarm may nevertheless over-cluster environmental molecular data (i.e., produce too few OTUs) if there is no or low variability between species (Mahé et al. 2017). Swarm may also under-cluster (i.e., produce too many OTUs) if the sequencing depth was too shallow (Mahé et al. 2015), or if there is not concerted evolution among multiple genomic copies of the targeted gene region, but the extent of such intra-cellular polymorphisms is debated (Brown et al. 2015, Decelle et al. 2014).

Together our sampling and molecular data did show that protists in the lowland Neotropical rainforests exhibit biogeographical patterns. Here we persistently found—for the first time—these patterns at the OTU-and phylogenetic-diversity levels, using taxa-area and distance-decay relationships, with presence-absence and abundance-based measures, on three large protist taxa as well as their subtaxa. Previous studies aimed at analyzing the biogeographies of protists inhabiting temperate soils only evaluated one or a few of these aspects (Bahram et al. 2016, Bates et al. 2013, Fernández et al. 2016, Lara et al. 2016, Venter et al. 2017). We additionally evaluated two null hypotheses in order to reject the random origin of the observed biogeographic patterns. Low taxa-area slopes were previously used to argue that protists and other microbes lacked biogeographies (Azovsky 2002, Finlay 2002, Horner-Devine et al. 2004). We showed here, however, that there could be significant biogeographic structures even with low taxa-area and distance-decay slopes, as long as the slopes were steeper than the slopes produced by the null models in which either the sample locations or the phylogenetic placements were randomly shuffled.

The biogeographical patterns of high alpha diversity within forests and low beta diversity between forests were persistent for both the parasitic Apicomplexa and the free-living Cercozoa and Ciliophora, even though they have different ecological functions and different dispersal modes. The apicomplexan patterns can be explained by them mirroring the patterns of their animal hosts: arthropod biogeographic patterns, at least for the herbivores, are themselves partially driven by their host plants (Basset et al. 2015, Novotny et al. 2007). Apicomplexan patterns should follow the animals if they are largely host-specific, although little is known of their biology in the Neotropics except for some Haemospororida, such as *Plasmodium* (Fecchio et al. 2017, Svensson-Coelho et al. 2016), which accounted for only a small fraction of Apicomplexa here. The cercozoan and ciliophoran patterns within forests suggest that their OTU diversity is driven by environmental filtering, while their phylogenetic diversity is structured by spatially-related processes, at least partially; similar patterns were also observed for Neotropical trees within forests (Swenson et al. 2013). Between forests, both of these free-living taxa showed spatial patterns of OTU diversity suggesting that their biogeographies are partially structured by dispersal limitations, while only Ciliophora exhibited spatial phylogenetic pattern highlighting the influence of evolutionary history processes on their biogeographic structure.

These patterns of taxa-area and distance-decay relationships for the microbial protists were likewise found in tropical macro-organismic animals and plants (Basset et al. 2012, Condit et al. 2002, Novotny et al. 2007, Pitman et al. 1999): there was high alpha diversity within forests, but low beta diversity between forests. Our study therefore demonstrates that many of the same general processes partially structure the biogeography of macro- and micro-organismal eukaroytes in Neotropical lowland rainforests. Unlike the animals and plants, though, for the protists we did not observe distance-decay patterns within forests; this difference could be due to the extremely high diversity of protists at each sample location, or due to the sampling scale (such patterns could possibly be observed for protists at the cm² or m² scale). These biogeographical observations at the OTU levels are pertinent to the problem of estimating the local and regional diversity of eukaryotes (Mora et al. 2011). As with estimating the diversity of tropical arthropods (Novotny et al. 2002, Novotny et al. 2007), this low regional turnover means that it is not simply a matter of discovering all protist species within a forest and then extrapolating that number by the total area of the Neotropics. And as with arthropods (Basset et al. 2012, Basset et al. 2015), the problem remaining for the protists is how to effectively estimate their total diversity within a small area because of the low local similarity in OTUs between samples (i.e., how many samples must be taken to estimate the total number of protists in just the soils of a forest, as well as those that are inhabiting the tree canopy and other microhabitats), as well as determining the most appropriate scale to describe local biogeographic structures.

## ACKNOWLEDGMENTS

We thank Jordan Mayor and Tobias Siemensmeyer for help in collecting. Funding came from the Deutsche Forschungsgemeinschaft grants DU1319/1-1 and DU1319/1-2 to M.D.

---

**Figure S1.**
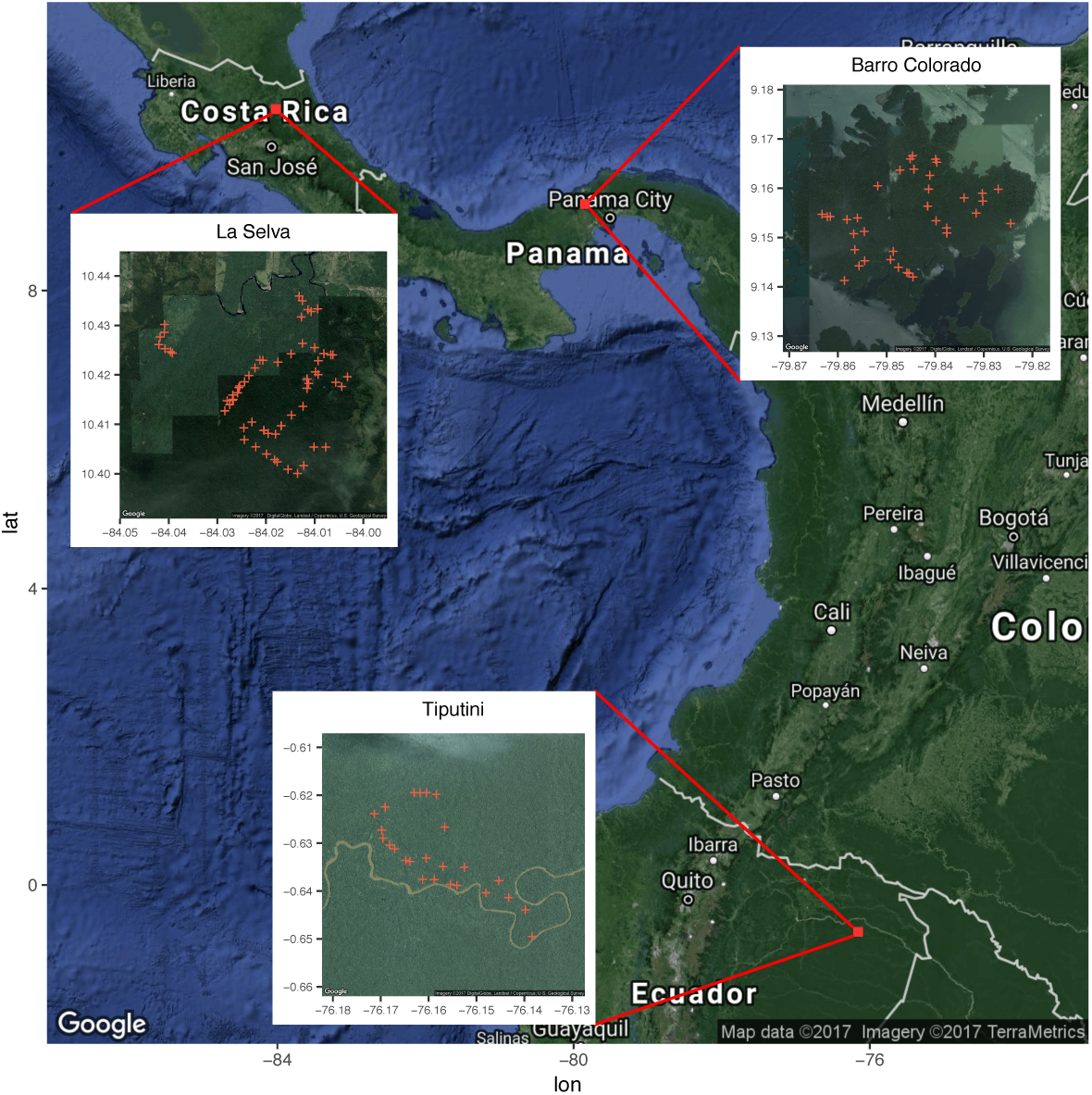
Map of the sampled Neotropical region with inset into sampled forests in Costa Rica (La Selva Biological Station), Panama (Barro Colorado Island) and Ecuador (Tiputini Biodiversity Station). Red crosses are individual sampling points.

**Figure S2.**
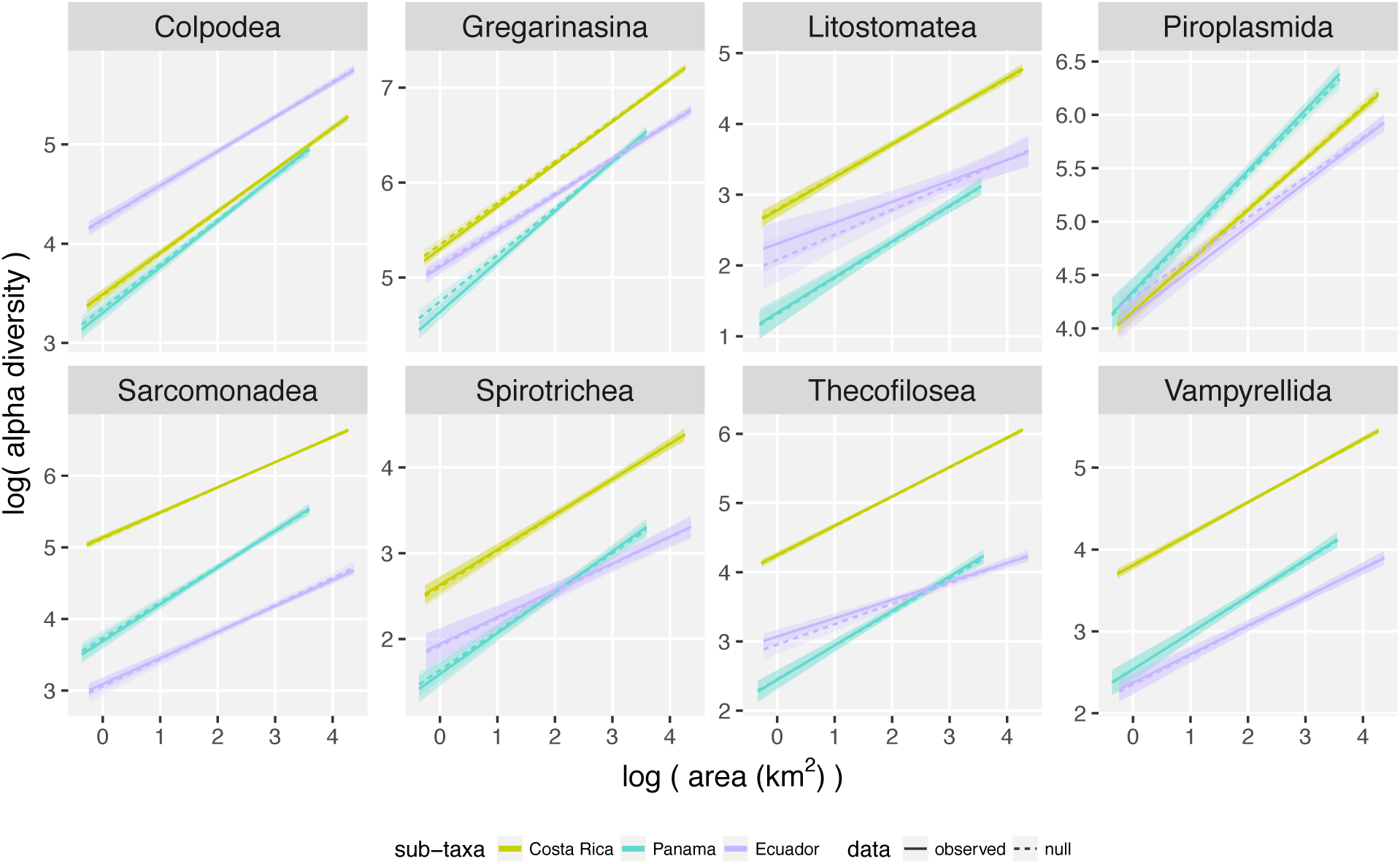
OTU richness based Taxa-Area Relationship (TAR) within forests for each sub-taxa. None of the slopes significantly differed from null model 1.

**Figure S3.**
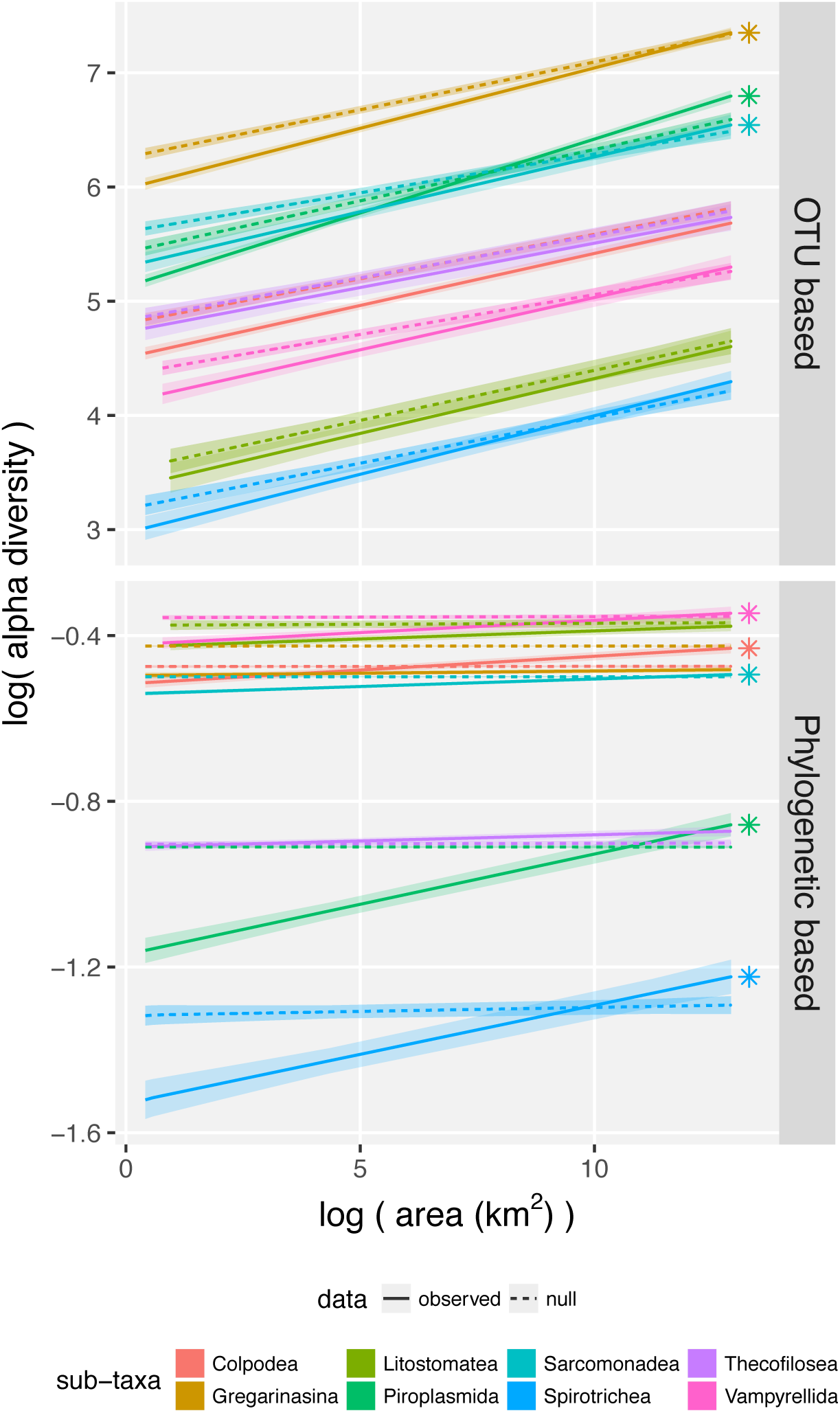
Taxa-Area Relationship (TAR) of sub-taxa across forests. Stars denoted significant steepest slopes than the null model. Null model 1 (samples random placement) was used for OTU based analyses and null model 2 (phylogenetic tree tips shuffling) was used for phylogenetic based analyses.

**Figure S4.**
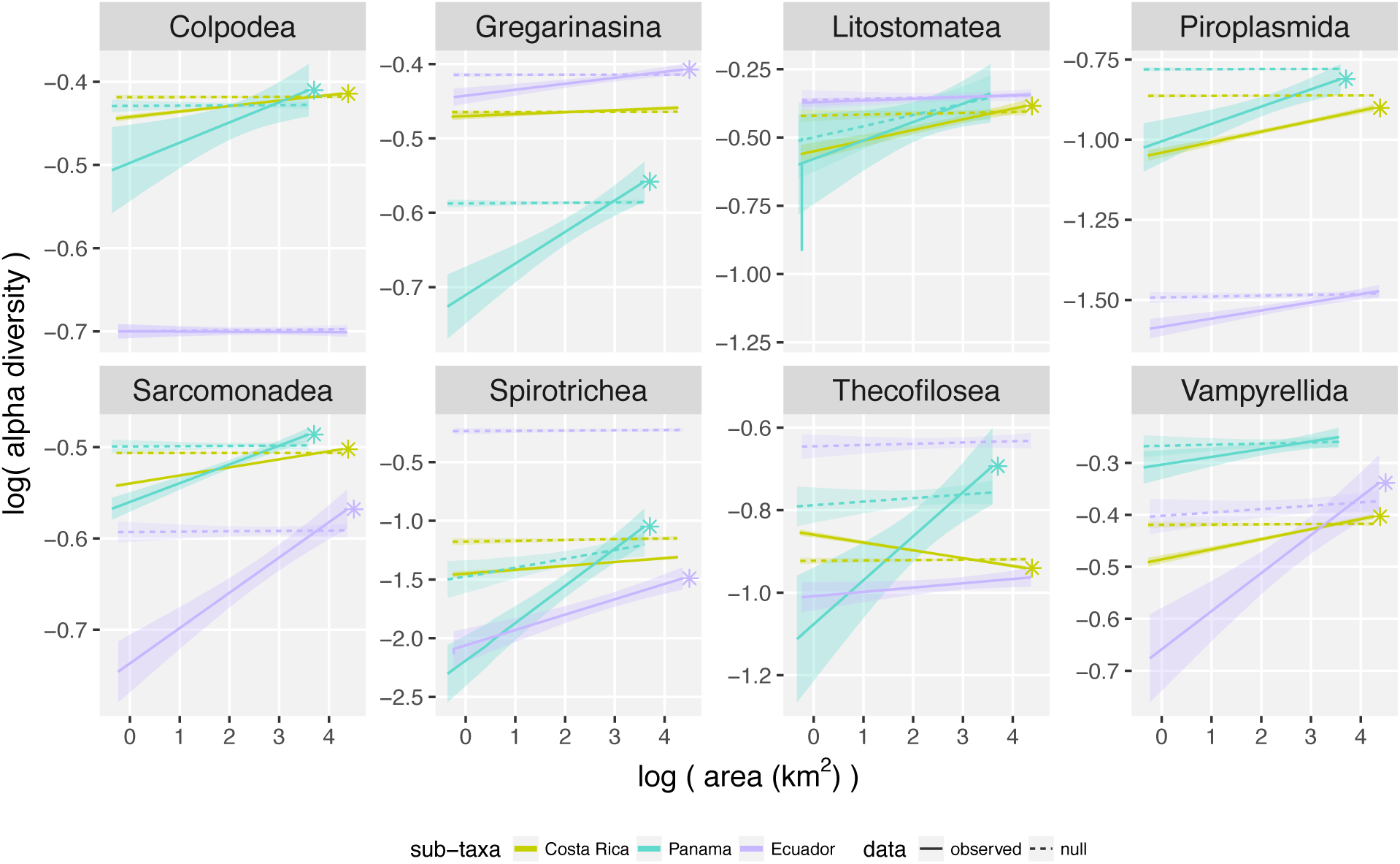
Phylogenetic variability based Taxa-Area Relationship (TAR) within forests for each sub-taxa. Stars denoted significant steepest slopes compared to null model 2.

**Figure S5.**
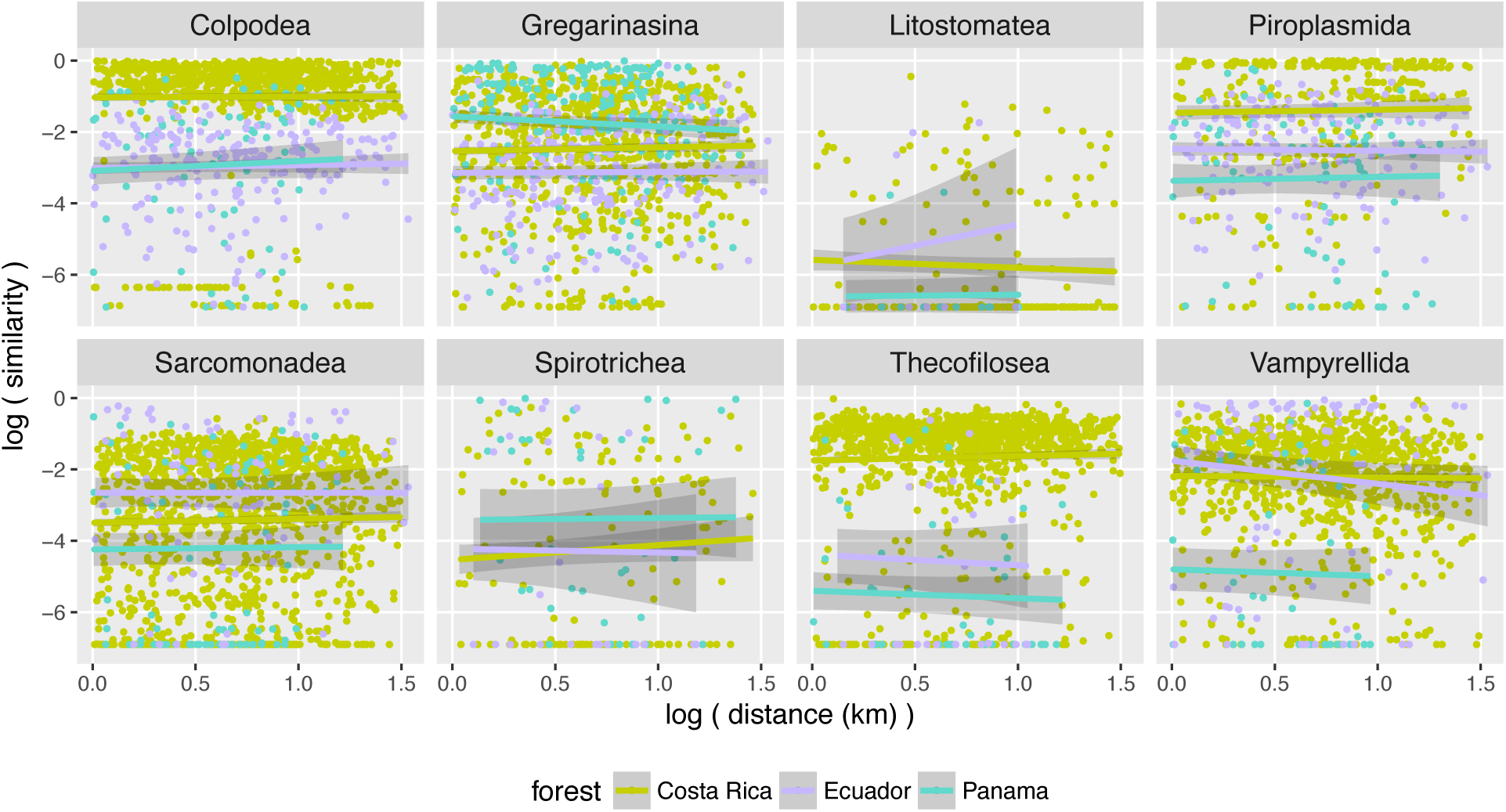
OTU relative abundance based Distance-Decay Relationship (DDR) of sub-taxa within forests. All fitted linear regression lines were not significant.

**Figure S6.**
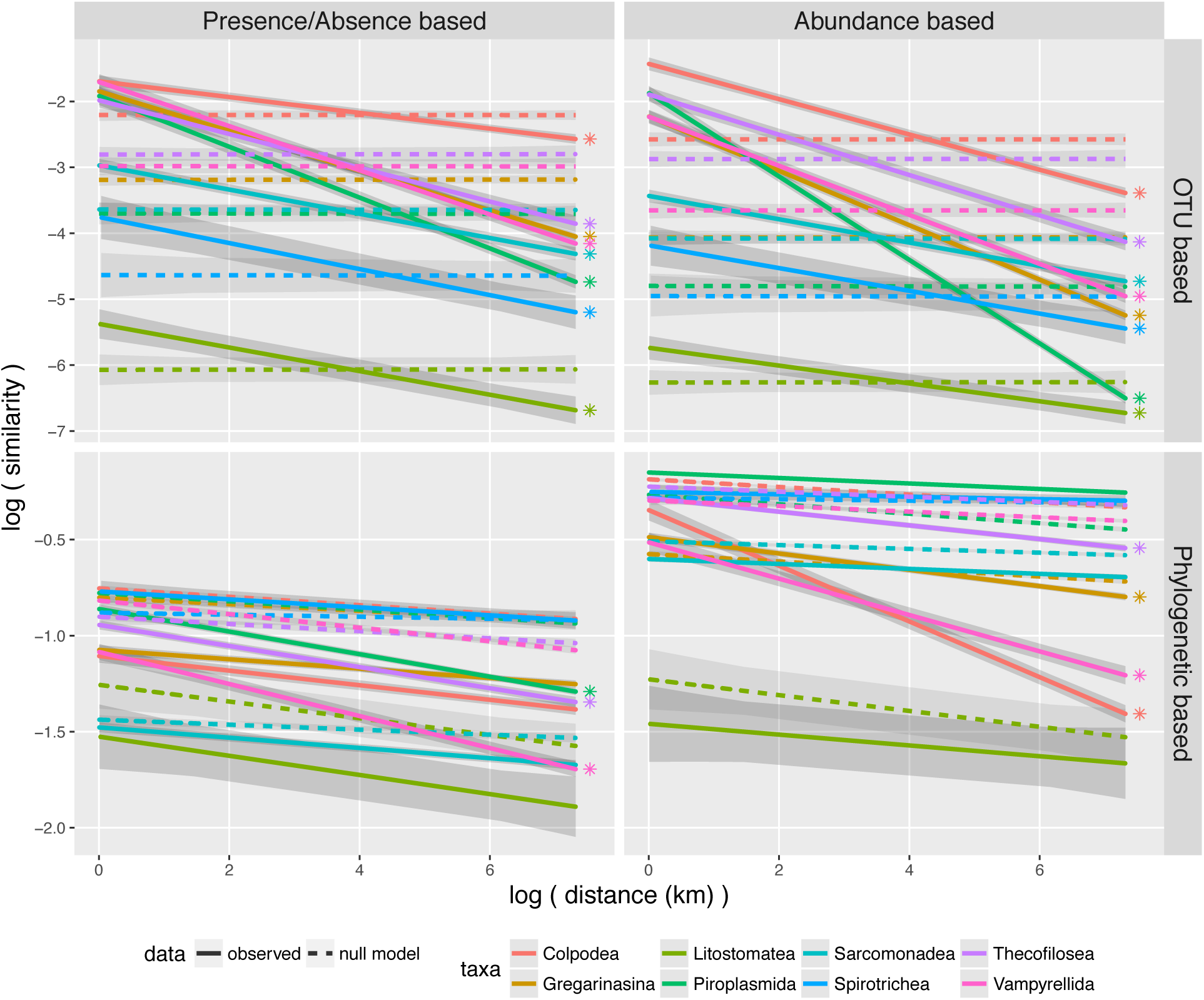
Distance-Decay Relationship (DDR) of sub-taxa across forests. A value of 1e-3 was added to similarity indexes to avoid infinite negative values in the log space. Stars denoted significant steeper slopes than null model. Sub-taxa OTU based DDR slopes were all significantly negative and steepest than the null model 1. Only Thecofilosea and Vampyrellida have significant steeper phylogenetic based DDR slopes than in the null model 2 for both presence/absence and abundance based indexes. Piroplasmida have significant steeper phylogenetic based DDR slopes than in the null model 2 for the presence-absence based index while this was true for Colpodea and Gregarinasina only with the abundance based index.

**Table S1.**
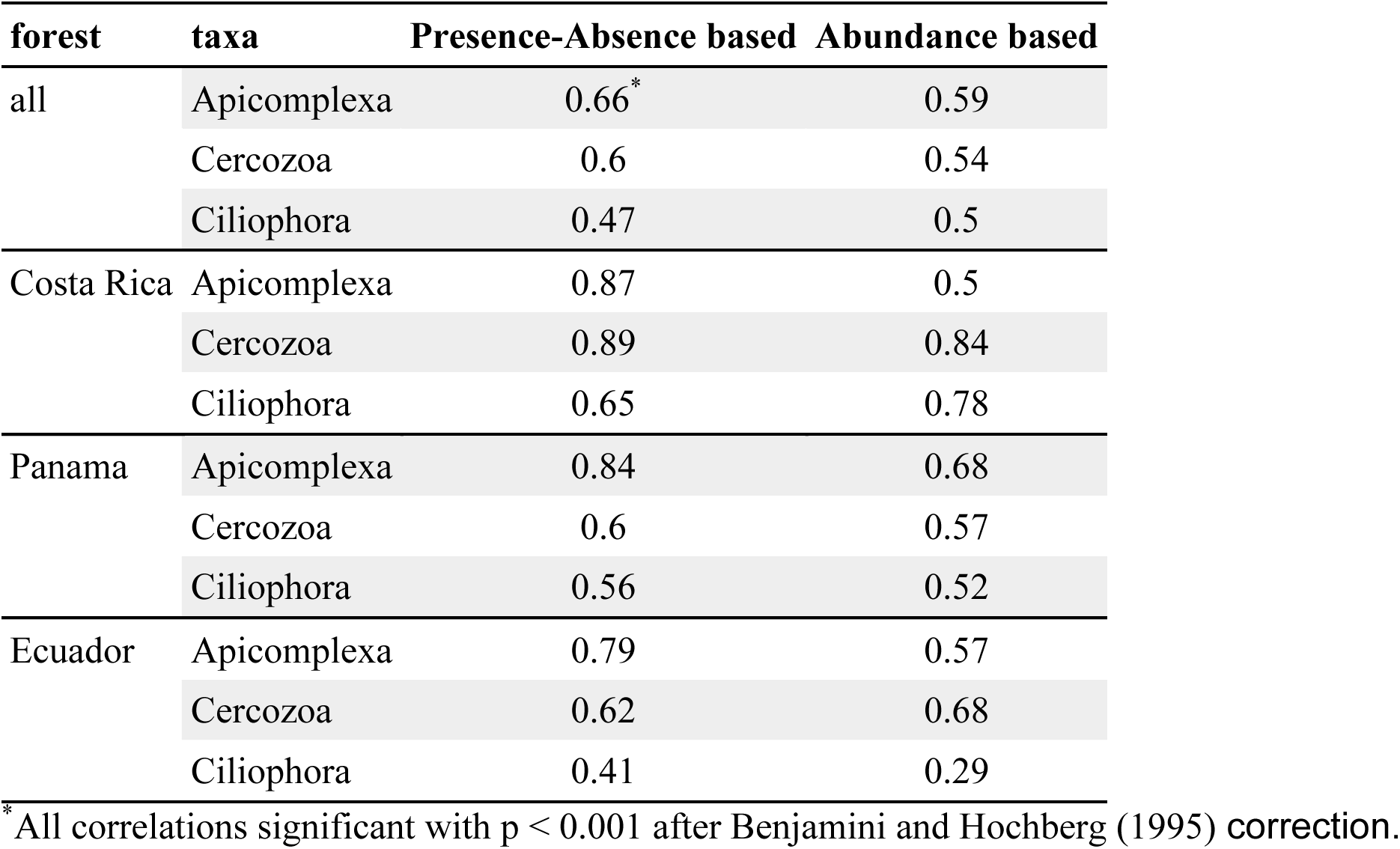
Spearman’s rho correlation coefficient calculated between OTU-and phylogenetic-based community similarity measures.

| forest | taxa | Presence-Absence based | Abundance based |
| --- | --- | --- | --- |
| all | Apicomplexa | 0.66* | 0.59 |
|  | Cercozoa | 0.6 | 0.54 |
|  | Ciliophora | 0.47 | 0.5 |
| Costa Rica | Apicomplexa | 0.87 | 0.5 |
|  | Cercozoa | 0.89 | 0.84 |
|  | Ciliophora | 0.65 | 0.78 |
| Panama | Apicomplexa | 0.84 | 0.68 |
|  | Cercozoa | 0.6 | 0.57 |
|  | Ciliophora | 0.56 | 0.52 |
| Ecuador | Apicomplexa | 0.79 | 0.57 |
|  | Cercozoa | 0.62 | 0.68 |
|  | Ciliophora | 0.41 | 0.29 |

\* All correlations significant with $p < 0.001$ after Benjamini and Hochberg (1995) correction.

